# Pathogens, endosymbionts, and blood-meal sources of host-seeking ticks in the fast-changing Maasai Mara wildlife ecosystem

**DOI:** 10.1101/2020.01.15.907568

**Authors:** Joseph Wang’ang’a Oundo, Jandouwe Villinger, Maamun Jeneby, George Ong’amo, Moses Yongo Otiende, Edward Edmond Makhulu, Ali Abdulahi Musa, Daniel Obado Ouso, Lillian Wambua

## Abstract

**Background:** The role of questing ticks in the epidemiology of tick-borne diseases in Kenya’s Maasai Mara National Reserve (MMNR), an ecosystem with intensified human-wildlife-livestock interactions, remains poorly understood. Therefore, we carried out a survey of the diversity of questing ticks, their blood-meal hosts, and tick-borne pathogens to understand potential effects to human and livestock health.

**Methods:** Questing ticks were collected by flagging and hand picks from vegetation in 25 localities and identified based on morphologic and molecular criteria. We used PCR with high-resolution melting (HRM) analysis, and sequencing to identify *Anaplasma, Babesia, Coxiella, Ehrlichia, Rickettsia*, and *Theileria* pathogen diversities and blood meals in 231 tick pools.

**Results:** A total of 1,465 host-seeking ticks were collected, including *Rhipicephalus appendiculatus* (n = 1,125), *Rhipicephalus pulchellus* (n = 6), *Rhipicephalus evertsi* (n = 5), *Amblyomma* cf. *gemma* (n = 178), *Amblyomma gemma* (n = 145), *Amblyomma variegatum* (n = 4), *Amblyomma* sp. (n = 1), and *Haemaphysalis leachi* (n = 1). Remnant blood-meals from humans, wildebeest, and African buffalo were detected in *Rh. appendiculatus*, goat in *Rh. evertsi*, sheep in *Am. gemma*, and cattle in *Am. variegatum. Rickettsia africae* was detected in *Am. gemma* (1/25 pools) that had blood-meal remnant from sheep and *Am. variegatum* (4/25 pools) that had fed on cattle. *Rickettsia* spp. were found in *Am. gemma* (4/25 pools) and *Rh. evertsi* (1/4 pools). *Anaplasma ovis* was detected in *Rh. appendiculatus* (1/172 pools) and *Rh. evertsi* (1/4 pools), while *Anaplasma bovis* was detected in *Rh. appendiculatus* (1/172 pools). *Theileria parva* was detected in *Rh. appendiculatus* (27/172 pools). *Babesia, Ehrlichia* and *Coxiella* pathogens were not found in any ticks. Unexpectedly, diverse *Coxiella* sp. endosymbionts were detected in all tick genera (174/231 pools).

**Conclusions:** The data shows that ticks from the rapidly-changing MMNR are infected with zoonotic *R.africae* and unclassified *Rickettsia* spp, demonstrating the persistent risk of African tick-bite fever and other and Spotted Fever Group rickettsioses to local dwellers and visitors to the Maasai Mara ecosystem. Protozoan pathogens that may pose risk to livestock production were also identified. We also highlight possible existence of morphotypic variants of *Amblyomma* species, based on the observation of *Ambyomma* cf. *gemma*, which may be potential human parasites or emergent disease vectors. Our findings also demonstrate that questing ticks in this ecosystem have dynamic vertebrate blood sources including humans, wildlife and domestic animals, which may amplify transmission of tickborne zoonoses and livestock diseases. Further studies are needed to determine the role of *Coxiella* endosymbionts in tick physiology and vector competence.

## 1. Introduction

Wildlife ecosystems are known to be hotspots for a range of emerging diseases threatening human and livestock health [1–3]. The ecology of tick-borne pathogens (TBPs) is complex, often involving wildlife, domestic animals, and humans that not only provide blood-meals to maintain the tick populations, but also serve as reservoirs and/or amplifiers of different TBPs [4,5]. The majority of emerging pathogens are maintained asymptomatically by wildlife and are transmitted to humans and livestock by vectors such as ticks and mosquitoes. An upsurge of emerging tick-borne zoonoses has been witnessed globally in the recent decades, such as Lyme borreliosis, which affect humans in several developed countries [6], while the burden some of the “old” tick-borne diseases such as East Coast fever (ECF), have persisted [7]. Therefore, the study and control of TBPs demands for a ‘One Health’ approach, requiring knowledge of the tick species, their host feeding preferences, habitat, and range [5].

The emergence and expansion of TBDs is increasingly linked to changes in the physical environment [8,9]. It has been observed that ecosystems undergoing drastic changes (such as rapid vegetation cover degradation, changes in climate and land-use patterns) are likely to become “pathogenic landscapes” due to the increased connectivity and probability of contact between vectors and their animal and human hosts [10]. The Maasai Mara ecosystem in south-western Kenya, represents one such fast-changing environment. While this ecosystem has been long recognized as a biodiversity hub and the home of the spectacular wildebeest migration termed “*the 7*^*th*^ *wonder of the world*”, it has faced severe threats and challenges in the last three decades driven by drastic changes in land use [11–13]. Significant land fragmentation has occurred in the Maasai Mara to accommodate an increasing number of conservancies, tourist lodges, human settlements, and agricultural developments [14]. Multiple land uses, such as pastoralism, commercial ranching, camping, tourism, and illegal grazing are being practiced concurrently, which favor the convergence of humans and domestic animals with wild animals and increase the risk of pathogen transmission [14–17]. Therefore, we hypothesized that the Maasai Mara National Reserve (MMNR) may serve as a model to study interactions between ticks, pathogens, and their vertebrate hosts in a fast-changing environment. We based our study on questing ticks because this is the active stage of ixodid ticks seeking vertebrate hosts for blood-meals in which they pick up new pathogens and transmit them from previous blood-meals to new hosts, including humans and livestock.

## 2. Materials and methods

### 2.1. Study area

The MMNR lies within Narok County in southwestern Kenya (Fig 1) and is contiguous with the Serengeti National Park in northern Tanzania. The MMNR supports a high diversity of large and small mammals and is globally famous for the annual wildebeest migration involving 1.3 million wildebeest, 200,000 zebras and hundreds of thousands of Thomson’s gazelles, topi, and elands [14]. This great migration into the Maasai Mara begins in July when these animals migrate from the Serengeti plains and ends in October when they migrate back to the Serengeti. The MMNR also includes tourist lodges, hotels, conservancies and commercial ranches, which act as wildlife dispersal areas. There are also settlements inhabited by local indigenous Maasai tribe whose livelihoods are dependent on livestock, mainly cattle, goats and sheep. Ethical clearance for this research in protected areas was sought from and approved by the Kenya Wildlife Service (KWS) Research Authorization committee.

**Fig 1.**
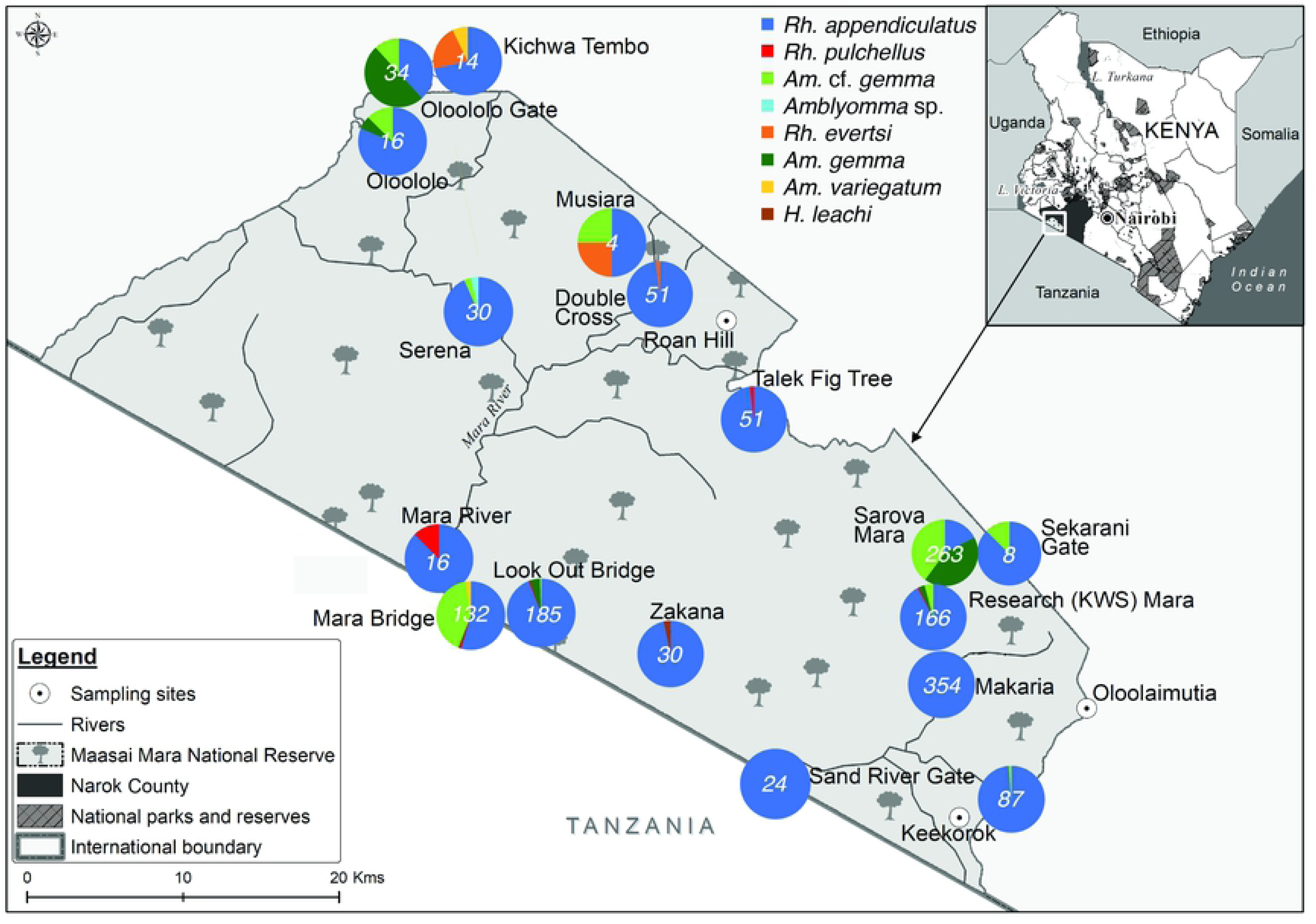
Distribution and abundance of tick species across various study sites of the Maasai Mara National Reserve.

### 2.2. Tick collection and identification

Questing ticks were collected from 25 sites in the MMNR during the great wildebeest migration in June – July of 2016 (Fig 1). Within each sampling site, we established survey plots measuring 100 m × 100 m comprising vertebrate resting areas, burrows, host routes and watering holes. Questing ticks were sampled within these plots between 10am and 5pm using a combination of flagging and picking from vegetation with gloved hands and a pair of forceps [18]. Flagging was carried out by slowly dragging a 1m^2^ white cotton cloth over the vegetation along 100-m transects. Ticks attached to the cloth were collected using forceps after each 10-m drag, put in sterile labelled tubes and frozen on dry ice while in the field, before being carried in liquid nitrogen to the lab in International Centre of Insect Physiology and Ecology (*icipe*), Nairobi. Once in the lab, adult ticks were identified to genus and/or species level under a microscope (Stemi 2000-C, Zeiss, Oberkochen, Germany) based on morphological keys developed by Walker et al. [19]. Immature stages were identified to genus level against larvae and nymphs of conspecific adults of *Rhipicephalus* and *Amblyomma* laboratory colonies maintained in *icipe*’s tick unit and further confirmed by molecular analyses. To prevent contamination with exogenous DNA, sterile petri-dishes, gloves, forceps, and gloves were used while handling the ticks. After morphological identification, ticks were pooled according to sampling site, species, and sex into groups of 1-11 adults, 1-20 nymphs, and 1-25 larvae.

### 2.3. DNA extraction from tick pools

Tick pools were homogenized in 1.5 ml Eppendorf tubes using zirconium oxide beads (Glen Mills, Clifton, NJ, USA) as previously described [20]. Genomic DNA was extracted from the homogenates using a protein precipitation method. Briefly, 300 μl of sterile cell lysis buffer (10 mM Tris-HCL [pH 8.0], 5 mM EDTA, 0.5% SDS) was added to the homogenate. The lysate was then incubated at 65°C for 60 minutes, followed by addition of 100 μl of protein precipitation solution (8 M ammonium acetate, 1 mM EDTA). The tubes were vortexed for 30 seconds and centrifuged for 10 min at 14,000 x g. The supernatants were aspirated and 300 μl of isopropanol was added to precipitate the DNA. Tubes were vortexed for 30 seconds, and the DNA was pelleted by centrifugation for 30 minutes at 14,000 x g. The DNA pellets were washed with 300 μl ice-cold 70% ethanol and air-dried for 12 hours before suspension in 50 μl of distilled deionized water. The quality and quantity of the extracted DNA samples was measured using a Nanodrop ND-1000 spectrophotometer (Thermo Scientific, Waltham, MA, USA) and diluted to 50 ng/μl for PCR. All DNA extracts were stored at −20°C until further use.

### 2.4. Molecular analyses of tick species

For molecular confirmation of species and genetic diversity, cytochrome oxidase 1 (CO1), 16S ribosomal ribonucleic acid (rRNA), and internal transcribed spacer 2 (ITS-2) gene markers were amplified by PCR from tick genomic DNA. The PCRs were performed in 10 μl reaction volumes that included 100 ng of DNA template, 1X HOT FIREPol® Blend Master Mix (Solis Biodyne, Estonia), 500 nM of each primer (S1 Table) and 5 μl PCR grade water. The following thermocycling conditions were used: Initial denaturation at 95°C for 15 min followed by 35 cycles of denaturation at 95°C for 20 s, annealing at 55°C (16S rRNA and CO1) and 65°C (ITS 2) for 30 s and extension at 72°C for 1 min, and a final extension at 72°C for 5 min. A no-template control with ddH_2_O in place of DNA was included in each run. PCR products were purified for sequencing using ExoSAP-IT Enzymatic PCR Product Clean-Up Kit (USB Corporation, Cleveland, OH, USA) according to the manufacturer’s instructions and sent to Macrogen (Netherlands) for capillary sequencing.

### 2.5. Screening of tick-borne pathogen DNA by PCR and high-resolution melting (PCR-HRM) analysis

The tick DNA samples were further tested by PCR for the presence of pathogens belonging to the genera *Anaplasma, Coxiella, Ehrlichia, Rickettsia, Theileria*, and *Babesia* using genus-specific primers (S1 Table). The procedure entailed touch-down PCR amplifications followed by melting of the amplicons in an HRM capable thermal cycler (Qiagen, Germany). The assays were performed in 10-μl reaction volumes, containing final concentration of 1x HOT FIREPol EvaGreen HRM mix (no ROX) (Solis BioDyne), 500 nM of the respective forward and reverse primers (S1 Table), 100 ng of template DNA and 5 μl PCR grade water. DNA samples of *Anaplasma phagocytophilum, Ehrlichia ruminantium*, and *R. africae* from an earlier study [21] were used as positive controls and no-template controls were also included. The touch-down PCR thermocycling conditions included an initial denaturation at 95°C for 15 min, followed by 10 cycles of amplification including denaturation at 94°C for 20 s; annealing for 25 s at 63.5-53.5°C (decreasing by 1°C per cycle), and 72°C for 30 s, followed by 25 cycles with a constant annealing temperature of 50.5°C and a final extension step at 72°C for 7 min. Cycling conditions described by Fard and Khalili [22] were used for *C. burnetii-*specific primers. An HRM step was thereafter performed in which amplicons were gradually melted from 75-90°C with 0.1°C increments every 2 s. Melting profiles were visualized within the Rotor-Gene Q Software v.2.1.0 (Build 9). Representative amplicons associated with each unique HRM profile were purified for sequencing.

### 2.6. Analysis of tick blood-meal remnants by PCR-HRM

We investigated the vertebrate sources of tick blood-meals following established protocols [23,24]. Briefly, genomic DNA from individual tick pools were analyzed by PCR amplification of vertebrate cytochrome b (*cyt b*) and 16S ribosomal (r) RNA genes using primers listed in S1 Table. PCR reactions were set-up with similar reaction volumes and components as already described for pathogen PCR-HRM. DNA extracts from voucher wildlife specimens (obtained from the Kenya Wildlife Service) and livestock species listed here were included as positive controls: Blue wildebeest (*Connochaetes taurinus*), giraffe (*Giraffa camelopardalis*), impala (*Aepyceros melampus*), buffalo (*Syncerus caffer*), warthog (*Phacochoerus africanus*), Grant’s gazelle (*Nanger granti*), hartebeest (*Alcelaphus buselaphus*), waterbuck (*Kobus ellipsiprymnus*), plain’s zebra (*Equus quagga*), Kirk’s dik-dik (*Madoqua kirkii*), Sable antelope (*Hippotagus niger*), lion (*Panthera leo*), cattle (*Bos taurus*), sheep (*Ovis aries*), and goat (*Capra hircus*). The amplicons with unique melt curves were purified for confirmation by Sanger sequencing.

### 2.7. Genetic and phylogenetic analyses

Using the MAFFT plugin in Geneious software version 11.1.4 (created by Biomatters) [25], all study nucleotide sequences were edited and aligned with related sequences identified by querying in the GenBank nr database using the Basic Local Alignment Search Tool (www.ncbi.nlm.nih.gov/BLAST/). The aligned DNA sequences were used to construct maximum likelihood phylogenetic trees using PHYML v.3.0 [26]. The phylogenies employed the Akaike information criterion [27] for automatic model selection and tree topologies were estimated using nearest neighbor interchange (NNI) improvements over 1,000 bootstrap replicates. Phylogenetic trees were visualized using FigTree v1.4.2.

## 3. Results

### 3.1. Tick species diversity

We collected a total of 1,465 questing ticks across the MMNR including 1137 adults, 97 nymphs, and 231 larvae. According to morphological and genetic analysis, *Rhipicephalus appendiculatus* comprised the highest proportion of the ticks sampled (n = 1125, 76.79%). Other species included *Rhipicephalus pulchellus* (n = 6, 0.41%), *Rhipicephalus evertsi* (n = 5, 0.34%) *Amblyomma gemma* (n = 145, 9.90%), *Amblyomma variegatum* (n = 4, 0.27%) and *Haemaphysalis leachi* (n = 1, 0.07%) (Fig 1; Table 1). Additionally, we identified a proportion of *Amblyomma* ticks (n = 178, 12.15%), which were morphologically similar to *Am. gemma*, except that the joining of the posteromedian stripe to the falciform stripe was incomplete in these species, relative to *Am. gemma* in which the two stripes are fully joined in reference images (Fig 2; [19]). Nevertheless, the ornamentation of their mesial and lateral median areas on the scutum was similar to that of *Am. gemma*, and they exhibited partial enameling of the festoons with the central festoon being dark as in *Am. gemma* (Fig 2; [19]). Hence, for purposes of this work, these ticks were termed as *Amblyomma* cf. *gemma*. Further genetic analysis of the adult and immature stages *Am.* cf. *gemma* based on CO1, 16S rRNA and ITS-2 was done, but only the latter marker yielded amplicons. The ITS-2 sequences of the *Am.*cf. *gemma* clustered together with *Am. gemma, Am. hebraeum, Amblyomma eburneum* and *Amblyomma variegatum* (Fig 3). In addition, one adult *Amblyomma* male tick which could not be morphologically identified to species level (Fig 2) but had an ITS-2 sequence with 99% identity to that of an *Amblyomma* sp. removed from nostril of a traveler who had visited Lope National Park in Gabon (Fig 3; [28]) was identified. Hence, for purposes of this work, this tick was termed as *Amblyomma* sp. (Fig 2). The ITS-2 sequences of eight *Rh. appendiculatus* nymphs were 100% identical to reference *Rh. appendiculatus* sequences (Fig 3).

**Table 1.** Ticks species, tick-borne pathogens and endosymbionts detected in ticks from this study.

| Tick species | Number of ticks |  |  |  |  | Number of tick pools | Number of pathogen and endosymbiont positive adult tick pools (%) |  |  |  |  |  |  |  |  |
| --- | --- | --- | --- | --- | --- | --- | --- | --- | --- | --- | --- | --- | --- | --- | --- |
|  | Total number (percentage) | L | N | AM | AF |  | *Rickettsia* spp. | *R. africae* | *A. bovis* | *A. ovis* | *T. parva* | *Babesia* spp. | *C. burnetii* | *Ehrlichia* spp. | *Coxiella* sp. endosymbionts |
| *Rhipicephalus appendiculatus* | 1,125 (76.79%) | 30 | 8 | 470 | 617 | 172 | 0 | 0 | 1 (0.58%) | 1 (0.58%) | 27 (15.70%) | 0 | 0 | 0 | 138 (80.23%) |
| *Rhipicephalus pulchellus* | 6 (0.41%) | 0 | 0 | 2 | 4 | 5 | 0 | 0 | 0 | 0 | 0 | 0 | 0 | 0 | 5 (100%) |
| *Rhipicephalus evertsi* | 5 (0.34%) | 0 | 0 | 2 | 3 | 4 | 1 (25.00%) | 0 | 0 | 1 (25.00%) | 0 | 0 | 0 | 0 | 1 (25.00%) |
| *Amblyomma* cf. *gemma* | 178 (12.15%) | 91 | 76 | 11 | 0 | 20 | 0 | 0 | 0 | 0 | 0 | 0 | 0 | 0 | 9 (45.00%) |
| *Amblyomma gemma* | 145 (9.90%) | 110 | 13 | 10 | 12 | 25 | 3 (12.00%) | 1 (4.00%) | 0 | 0 | 0 | 0 | 0 | 0 | 18 (72.00%) |
| *Amblyomma variegatum* | 4 (0.27%) | 0 | 0 | 1 | 3 | 3 | 0 | 1 (33.33%) | 0 | 0 | 0 | 0 | 0 | 0 | 2 (66.67%) |
| *Amblyomma* sp. | 1 (0.07%) | 0 | 0 | 1 | 0 | 1 | 0 | 0 | 0 | 0 | 0 | 0 | 0 | 0 | 0 (0.00%) |
| *Haemaphysalis leachi* | 1 (0.07%) | 0 | 0 | 0 | 1 | 1 | 0 | 0 | 0 | 0 | 0 | 0 | 0 | 0 | 1 (100%) |
| **Total** | **1,465** | **231** | **97** | **497** | **640** | **231** | **4 (1.73%)** | **2 (0.87%)** | **1 (0.43%)** | **2 (0.87%)** | **27 (11.69%)** | **0** | **0** | **0** | **174 (75.32%)** |

**Abbreviations:** L - larvae, N – nymphs, AM – adult males, AF – adult females

**Fig 2.**
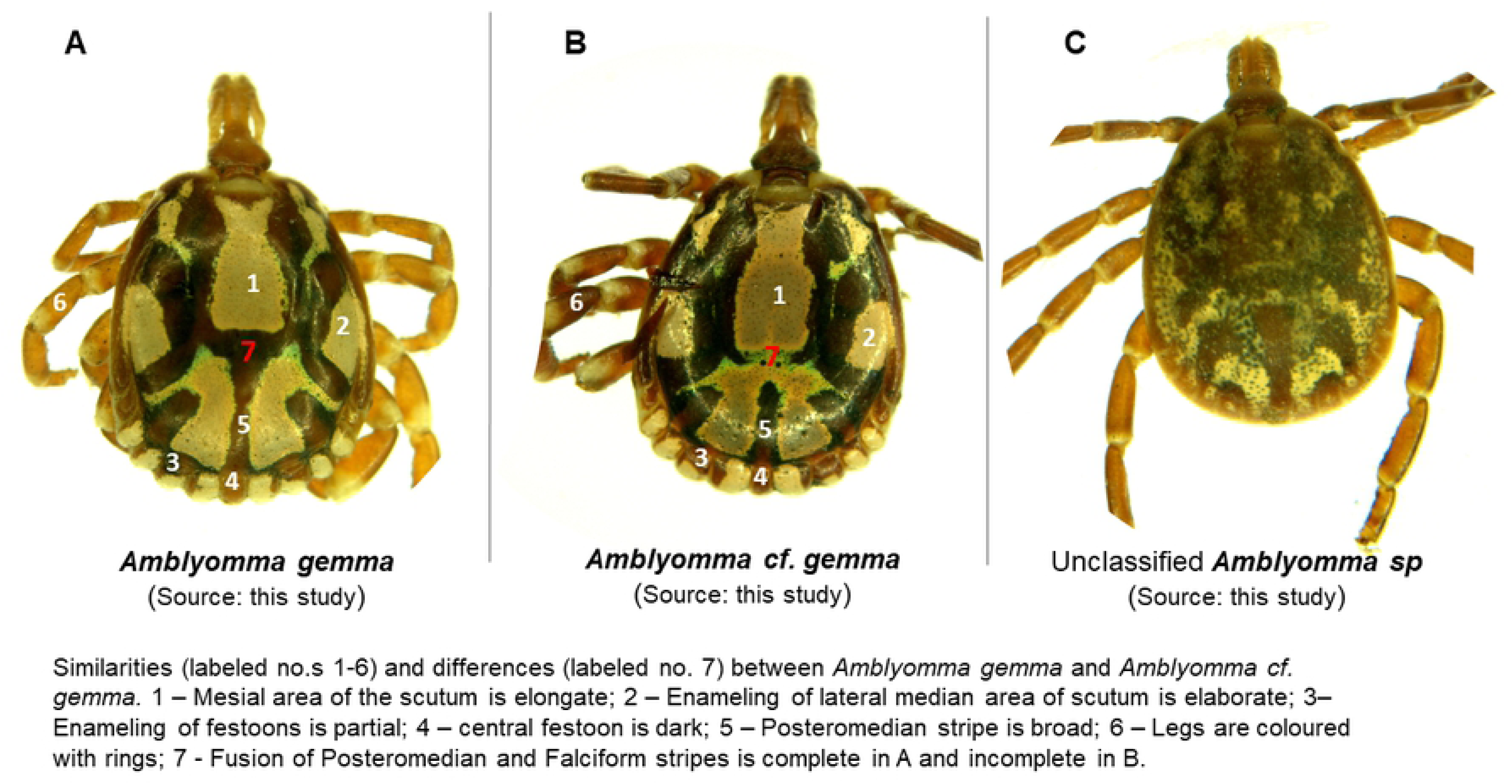
Comparison of selected *Amblyomma* species from this study: *Amblyomma gemma* and *Amblyomma* cf. *gemma* ticks are depicted on panels **A** and **B** while an unclassified *Amblyomma* tick is shown in panel **C**. Similarities between *Amblyomma gemma* and *Amblyomma* cf. *gemma* are labelled in white and differences in red colors respectively.

**Fig 3.**
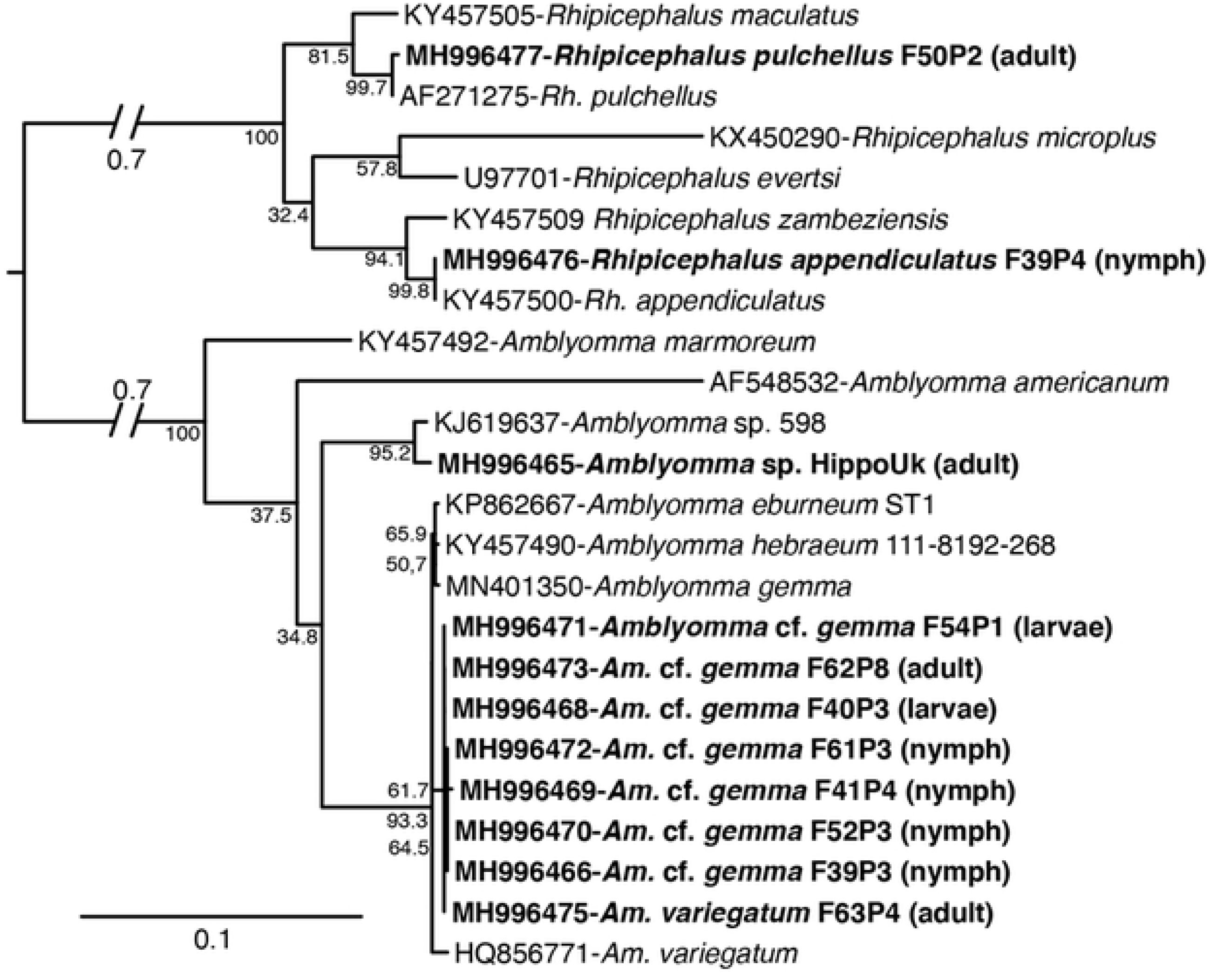
Maximum likelihood phylogenetic tree of tick ITS-2 gene sequences (739-1276 bp). GenBank accession numbers, species identifications, and isolates, and tick life stages (in brackets) are indicated for each sequence. Sequences from this study are indicated in bold letters. Bootstrap values at the major nodes are of percentage agreement among 1000 replicates. The branch length scale represents substitutions per site.

### 3.2. Tick-borne pathogens identified

A total of 231 pools of ticks were screened for *Anaplasma, Babesia, Coxiella, Ehrlichia, Rickettsia*, and *Theileria* pathogen infections, based on the pooling strategy depicted on Table 1. The pathogens detected are summarized in Fig 4 and Table 1. None of the pools showed any amplification for *Babesia* spp., *Ehrlichia* spp., or *Coxiella burnetii*.

**Fig 4.**
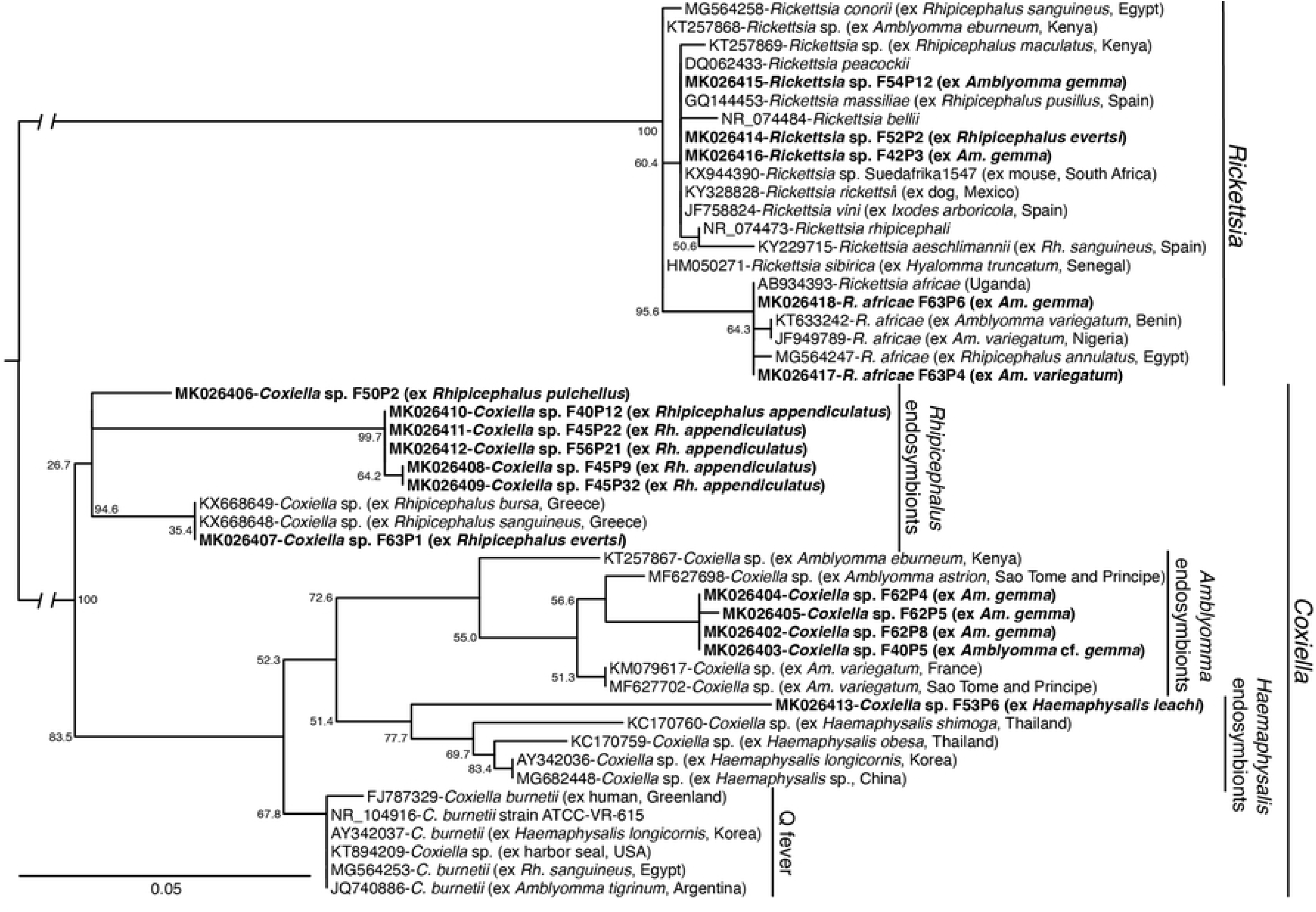
Maximum likelihood phylogenetic tree of *Rickettsia* and *Coxiella* 16S rRNA gene sequences (248-301 bp). GenBank accession numbers, species identifications, and isolates, with tick or vertebrate host species and country of origin in brackets, are indicated for each sequence. Sequences from this study are bolded. Bootstrap values at the major nodes are of percentage agreement among 1000 replicates. The branch length scale represents substitutions per site. The gaps indicated in the branches to the *Coxiella* and *Rickettsia* clades represent 0.10 substitutions per site.

Rickettsial DNA was amplified in six of the tick pools. Of these, *Rickettsia africae* 16S rDNA sequences sharing 100% identity with an isolate from Uganda were detected in two tick pools comprising of one adult *Am. gemma* female and one adult *Am. variegatum* male, both from Kichwa Tembo area of MNNR (Fig 4; Table 1; S2 Table). Additionally, DNA of unclassified *Rickettsia* spp. were amplified from one *Rh. evertsi* female tick and two *Am. gemma* (one male tick and one female) ticks. All but one of these rickettsial 16S rDNA sequences were 99-100% identical to *Rickettsia* sp. Suedafrika1547 accession KX944390 and clustered with other spotted fever group (SFG) rickettsiae, including *Rickettsia massiliae, Rickettsia rhipicephali, Rickettsia amblyommii*, and *Rickettsia raoultii* based on maximum likelihood phylogenetic analysis of a partial sequence of the 16S rRNA gene (Fig 4). On further attempts to classify the *Rickettsia* spp. to the species level using the ompB gene, amplicons from two pools of *Am. gemma* similarly showed 97% identity to *Rickettsia aeschlimannii, R. rhipicephali, R. massiliae* and *R. raoultii* (GenBank accessions MF002557, CP013133, KT835123, FN651773 respectively).

*Anaplasma* spp. 16S rDNA was amplified in two pools of *Rh. appendiculatus* and one pool of *Rh. evertsi* adult ticks (Table 1). The *Anaplasma* 16S rDNA sequences in one of the *Rh. appendiculatus* adult pools (comprising of seven male ticks from Double Cross area) was 100% identical to reference *Anaplasma bovis* (S2 Table). The *Anaplasma* 16S rRNA sequences from the other *Rh. appendiculatus* adult pool (comprising of seven female ticks sampled from Look Out Bridge area) and the *Rh. evertsi* adult pool (comprising of a female tick sampled from Kichwa Tembo area) were 100% identical to reference *Anaplasma ovis* (S2 Table).

*Theileria* spp. 18S rRNA sequences were amplified from 27 out of 172 pools of *Rh. appendiculatus* adult ticks collected from Makaria, Sarova Mara, Mara Bridge and Oloololo Gate regions of MMNR (Table 1; S2 Table). Sequences from representative samples of positive pools were 100 % identical to *T. parva* isolates detected in Zambia, Kenya, Uganda, and Tanzania (S2 Table).

### 3.3. Tick-borne endosymbionts identified

A total of 174 pools showed positive amplification for *Rickettsia* 16S rRNA but were negative for the *Rickettsia* ompB gene. Upon further sequencing and analysis, the amplicons revealed the presence of *Coxiella* sp. endosymbionts. These tick pools with *Coxiella* sp. endosymbionts yielded no amplification when screened with *C. burnetii*-specific primers. *Coxiella* sp. endosymbionts were detected in 80.23% of the *Rh. appendiculatus* pools (138/172), 100% of *Rh. pulchellus* pools (5/5,), 25% of the *Rh. evertsi* pools (1/4), 72% of *Am. gemma* pools (18/25), 45% of *Am.* cf. *gemma* pools (9/20), 66.67% *Am. variegatum* pools (2/3), and in the single *H. leachi* pool (Table 1). The endosymbionts were only absent in the single *Amblyomma* sp. tick reported in section 3.1 above. The *Coxiella* sp. endosymbionts were distributed across all sites in MMNR (S2 Table). Phylogenetic analysis demonstrated that these endosymbionts clustered in three host-specific clades associated with *Amblyomma, Rhipicephalus*, and *Haemaphysalis* ticks (Fig 4).

We further analyzed the tick pools with *Coxiella* sp. endosymbionts for co-infection with pathogens. The results showed that two pools of adult *Rh. appendiculatus* were co-infected with the aforementioned *A. ovis* and *A. bovis* (section 3.2). Additionally, of the 27 *Rh. appendiculatus* tick pools that were infected with *T. parva*, 70% (19/27) were co-infected with *Coxiella* sp. endosymbionts.

### 3.4. Vertebrate blood-meal sources in questing ticks

Only nine individual adult ticks had identifiable blood-meals (Table 2). Analysis of vertebrate *cyt b* sequences in these ticks revealed human blood-meals in one *Rh. appendiculatus*, and one *Am.* cf. *gemma*. Blue wildebeest (*Connochaetes taurinus*) blood-meals were detected in two *Rh. appendiculatus* ticks while African buffalo (*Syncerus caffer*) blood-meal was detected in one *Rh. appendiculatus* ticks. We also detected goat (*Capra hircus*) blood-meal in one *Rh. evertsi* tick. Amongst *Amblyomma* ticks, two *Am. gemma* ticks had blood-meals from sheep (*Ovis sp.*), while one *Am. variegatum* had a blood-meal from cattle (*Bos taurus*) (Table 2; Supplementary Fig. 1). The aforementioned *Rickettsia africae* infections (section 3.2) were detected in one of the *Am. gemma* ticks with a blood-meal from sheep and one of the *Am. variegatum* with a blood-meal from cattle.

**Table 2.** Summary of vertebrate sources of blood-meals in adult ticks detected by cytochrome b PCR and sequencing in questing ticks collected in this study.

| **Vertebrate source of blood-meal** | **Host tick species (proportion of tick species)** | **Sampling location** | **Submitted GenBank accessions** |
| --- | --- | --- | --- |
| Goat (*Capra hircus*) | *Rh. evertsi* (1/4) | Kichwa Tembo | MH997915 |
| Blue wildebeest (*Connochaetes taurinus*) | *Rh. appendiculatus* (2/172) | Sand River Gate and Zakaria | MH997918, MH997919 |
| African buffalo (*Syncerus caffer*) | *Rh. appendiculatus* (2/172) | Kichwa Tembo | MH997914 |
| Human (*Homo sapiens*) | *Rh. appendiculatus* (1/172) | Olololo Gate | MH997917 |
| Human (*H. sapiens*) | *Amblyomma* cf. *gemma* (1/20) | Olololo Gate | MH997916 |

## 4. Discussion

Concerns around the role of wildlife ecosystems as hotspots for a range of emerging diseases threatening human and livestock health have been rising, especially in areas where free-ranging wild animals regularly interact with domestic livestock and humans [1–3]. This study provides molecular evidence of presence of the zoonotic *R. africae*, uncharacterized *Rickettsia* spp. and veterinary pathogens (including *A. bovis, A. ovis*, and *T. parva*) in questing ticks collected from the MMNR. We also report that questing ticks in this wildlife ecosystem feed on humans, wildlife and domestic animals. Additionally, we report the presence of diverse species-specific *Coxiella* sp. endosymbionts in questing ticks in the MMNR. These findings are important to public and veterinary health strategies mitigating possible disease outbreaks in this fast-changing wildlife ecosystem in eastern Africa.

Diverse species of ticks were identified in this study, including *Amblyomma, Rhipicephalus*, and *Haemaphysalis* genera, which have previously been reported to occur in Maasai Mara region [29,30]. However, an interesting finding of this study, was the identification of an *Amblyomma* sp. comprising of 178 individuals with subtle degree of morphological variation relative to *Am. gemma*. These individuals were widely distributed across most study sites in this ecosystem, albeit not reported in previous studies in this locale or other regions of Kenya and for purposes of this study were provisionally termed *Am.* cf. *gemma*. Also, one individual, which was provisionally termed *Amblyomma* sp., had a distinct morphology, and was genetically closest by its ITS-2 sequence to a tick recovered from a traveler in Lope National Park in Gabon [28]. Although we could not conclusively deduce from the evidence in this study, whether these ticks were different species or morphotypes of known species, intraspecific morphological variations are known to occur in *Amblyomma* species [31] with studies elsewhere demonstrating phenotypic plasticity within this genus [32]. Morphotypic variation in *Am. gemma* has been reported by Walker and co-workers [19] where a minority of populations may exhibit incomplete fusion of the posteromedial and falciform stripes, as may be evident in this study. In the absence of careful morphological identification and given the geographical segregation of *Am. gemma* and *Am. hebraeum* across eastern and southern Africa, this subtle variation may potentially result in confusion between the two species of ticks. Nevertheless, the findings of this study, raise questions around the possible existence of species morphotypes or on-going morphological adaptation, which may be interesting in the context of this rapidly changing ecosystem. These results also highlight the need for morphological reassessment of *Amblyomma* species in this and other wildlife ecosystems in Africa coupled with extensive genetic studies using an array of both nuclear and mitochondrial markers.

Analysis of tickborne zoonotic pathogens in this study corroborate findings of previous studies in Kenya, which have demonstrated circulation of zoonotic SFG *Rickettsia* in ticks from various ecologies [21,33,34). The presence of *R. africae*, which causes a potentially fatal, but as yet neglected febrile illness, was first reported in this ecosystem in 2003 [29]. Since then, this pathogen has been highlighted as a threat to international travelers and local communities, yet in clinical settings in Kenya, no routine diagnosis for the pathogen in humans is done [35–40]. In this study, *R. africae* was detected in *Am. gemma* and *Am. variegatum* ticks, confirming the strong link between ticks of *Amblyomma* species and the epidemiology of *R. africae* in sub-Saharan Africa [34,35,41]. Further, *R. africae*-infected ticks were sampled from Kichwa Tembo area in MMNR, which is a wildlife-human interface dominated by tented camps and resorts. This may present a health hazard to local and international tourists visiting the reserve, further underpinning the need for continuous xenosurveillance and for clinicians in the Maasai Mara region and other wildlife ecosystems to include SFG rickettsiosis in differential diagnosis of febrile cases. Nevertheless, knowledge gaps on SGF rickettsiosis in Kenya persist, including the level of risk posed to humans by infected ticks, epidemiological evidence of the disease in human populations, and factors underlying infection and transmission dynamics of the pathogens by ticks and other vectors.

Unclassified *Rickettsia* spp. were also detected in *Rh. evertsi* and *Am. gemma* ticks, which may be potentially pathogenic based on positive amplification of the *ompB* gene, which encodes a membrane protein involved in adherence and invasion of host cells [42]. Previous reports have demonstrated high diversity of *Rickettsia* spp. and their potential vectors in Kenya [21,33,43]. Given that these reports have showed existence of novel species and uncharacterized variants of unknown pathogenicity, it will be important to undertake further studies of the unclassified *Rickettsia* spp. from this study, using combination of genetic markers such as *ompA, gltA, sca4*, 17kDa in addition to the *ompB* gene that was utilized in the present study. *Coxiella burnetii* was not detected in this study, augmenting prior findings of Ndeereh and co-workers [30], who also reported absence of *C. burnetii* in both ticks and animals in this ecosystem.

This study also found evidence of livestock pathogens in the ticks surveyed, key among which were *T. parva, A. bovis* and *A. ovis. Theileria parva* was the most prevalent livestock pathogen and was detected solely in *Rh. appendiculatus. Theileria parva* causes ECF, which remains the most economically important parasitic disease in cattle in eastern and southern Africa [19,44]. Therefore, our findings confirm that the distribution of *T. parva* is closely associated with *Rh. appendiculatus* ticks [44] and further that the *T. parva* in the MMNR were genetically identical to those found elsewhere in Kenya and within sub-Saharan Africa. These findings may imply that a similar control approach using the Infection and Treatment Method (ITM) with the live vaccine termed as the “*Muguga cocktail*” [45] can be applied in this setting in MNNR to control potential livestock losses from ECF. However, it has been demonstrated that proximity to buffaloes, the natural reservoir of *T. parva*, is associated with high *T. parva* diversity [46], which was not discernible from the current study. Nonetheless, the high prevalence of *T. parva* detected in *Rh. appendiculatus* ticks sampled in this study, indicates the need to comprehensively analyze the genotypes of *T. parva* isolates from ticks and cattle in in this ecosystem, relative to the proximity to buffalo niches and vaccination status.

Interestingly, tick species-specific *Coxiella* sp. endosymbionts were identified in 75.3% of adult ticks across the three genera sampled in this study, by sequencing amplicons obtained using the 16S rRNA primers for *Rickettsia* species described by Nijhof and co-workers [47]. Our results therefore suggest that these primers may have potential application in the study of *Coxiella* sp. endosymbionts in ticks, as none of the respective samples yielded positive PCR results when screened with *C. burnetii*-specific primers. While the current findings are interesting, there is need for subsequent studies to further characterize the *Coxiella* endosymbionts using housekeeping genes such as *fusA, rpsF*, and *rpsG* in addition to the 16S rRNA [48]. Previous investigations using the same assays as in this study, sequenced a *Coxiella* sp. endosymbiont from an *Am. eburneum* nymph, but not from other tick samples from the coastal region of Kenya [21] or from the Lake Victoria region [49]. This suggests that the prevalence of *Coxiella* sp. endosymbionts may depend on environmental factors. Their distinct association with particular species of ticks as shown in this study, prompts further investigations to establish whether or not the *Coxiella* sp. endosymbionts have potential roles in vector nutrition, reproductive fitness and vector competence of ticks in this ecosystem [50,51]. These findings also underpin the fact that PCR results of *Coxiella* infection in ticks must be interpreted with caution, especially if the amplified DNA products are not sequence [52–54].

The identification of blood-meal remnants in questing ticks from across the MMNR has lent insight into the feeding behavior of ticks and the risk of tick-borne diseases in this area. Although we amplified blood-meal remnants in only nine of the sampled adult questing ticks, diverse sources of vertebrate sources including goat, sheep, blue wildebeest, African buffalo, cattle, and humans were identified, indicating that the host-seeking patterns of ticks in the MMNR were dynamic. Nevertheless, the limited number of ticks with identifiable blood-meals was a limitation of the current study possibly because questing ticks can have their last blood-meal from their previous life stage up to one year before collection, which may have been degraded by digestive and hemolytic processes in the tick midgut [55,56]. It was further unclear whether the blood-meals identified in adult ticks were due to recent feeding or were remnants of nymphal feeding. Nevertheless, our finding that *Rh. appendiculatus* ticks had fed on humans, blue wildebeest, and African buffalo, not only confirms previous knowledge that *Rh. appendiculatus* infests a wide range of Bovidae [19,44], but also suggest a more diverse host-seeking pattern in involving humans. Although the known domestic hosts of adult *Rh. evertsi* are horses, donkeys, cattle, and sheep [19], we also detected a remnant blood-meal from goat. We also found that *Am. variegatum* from the MMNR had fed on cattle, which is consistent with previous report that all stages of this tick infest cattle, sheep, and goats [19]. The findings of blood-meals from domestic animals in a wildlife park setting also confirms the problem of encroachment of this wildlife ecosystem by humans and their livestock. The *Am.* cf. *gemma* ticks identified in this study had fed on humans and although no pathogens were identified in this species, it nevertheless highlights the species as a potential human parasite. We also detected *R. africae* infection in one *Am. gemma* that had fed on sheep and in one *Am. variegatum* that had fed on cattle. This suggests that sheep and cattle may be important in the epidemiology of African tick bite fever in the MMNR.

## 5. Conclusions

This study provides insights into the diversity of ticks, their microbes, and their blood-meal sources in the Maasai Mara ecosystem in Kenya, which is marked by rapid changes in land-use and considerable encroachment by humans and livestock. We report the observation of *Ambyomma cf. gemma*, and highlight possible existence of morphotypic variants of *Amblyomma* species, which may be potential human parasites and emergent disease vectors. We also demonstrate the presence and possible circulation of etiological agents of spotted group rickettsiosis that may pose serious constraints to human health in the MMNR, as well as anaplasmosis, and theileriosis that may impede livestock production. The results also highlight that tick species in the MMNR feed on humans as well as diverse wildlife and livestock including blue wildebeest, African buffalo, goat, sheep, and cattle. Our data also show that ticks from the MMNR harbor tick species-specific *Coxiella* spp. endosymbionts, highlighting the need for further studies to understand the role of these endosymbionts in tick physiology and vector competence.

## Availability of data from this study

Sequences obtained in this study have been deposited in the GenBank database under the following accession numbers: MH996465-MH996477 (ITS-2 sequences from ticks), MK026402-MK026413 (16S rRNA sequences of *Coxiella* sp. endosymbionts), MK026414-MK026418 (*Rickettsia* 16S rRNA), MK026419-MK026421 (*Anaplasma* 16S rRNA), MH997912-MH997913 (*Rickettsia* ompB genes), MH929321-MH929324 (*Theileria* 18S rRNA), MH997914-MH997919 (vertebrate *cyt b*).

## Acknowledgements

We acknowledge Antoinette Miyunga, Stephen Mwiu, Vasco Nyaga, Dennis Lemayian and Richard Bolo (all of KWS) and David Wainaina (*icipe*) for their help with the field surveys. We also acknowledge James Kabii (*icipe*) for his logistical assistance. We are also grateful for the technical assistance to Francis Matheka (Department of Veterinary Services in Kenya) in morphological identification of the ticks and Enock Mararo (*icipe*) in PCR-HRM analysis.

## Authors’ contributions

LW and JV designed the study. LW coordinated field activities. MYO coordinated the wildlife research permits for access to protected areas. LW, JV, MJ and DOO conducted the fieldwork. JWO, LW, EEM, and AAM performed laboratory analyses. JWO, LW, JV and GO conducted the data analysis. JWO wrote the first draft. All authors contributed to the manuscript editing and approved the final manuscript.

## Consent for publication

All authors consented and approved submission of this manuscript for publication.

## Supporting information

**S1 Fig. Alignment of short vertebrate 16S rRNA sequences of goat, sheep and cow blood-meals amplified from tick species in this study.** Sequences from this study are highlighted in (**bold**) against closest sequences available in GenBank. Colour code of nucleotides are depicted as Green = Thymine; Red = Adenine; Blue = Cytosine; Yellow = Guanine.

**S1 Table. PCR primer pairs used in this study**

**S2 Table: Detailed summary of geographical sources and sequence identities and Genbank accessions of tick-borne pathogens and endosymbionts detected in this study.**

## References

1. Jones KE, Patel NG, Levy MA, Storeygard A, Balk D, Gittleman JL, et al. Global trends in emerging infectious diseases. Nature. 2008 Feb;451(7181):990–993.

2. Siembieda JL, Kock RA, McCracken TA, Newman SH. The role of wildlife in transboundary animal diseases. Animal Health Research Reviews. 2011 Jun;12(01):95–111.

3. Wiethoelter AK, Beltrán-Alcrudo D, Kock R, Mor SM. Global trends in infectious diseases at the wildlife-livestock interface. Proceedings of the National Academy of Sciences of the United States of America. 2015 Aug;112(31):9662–7.

4. Parola P, Raoult D. Ticks and Tickborne Bacterial Diseases in Humans: An Emerging Infectious Threat. Clinical Infectious Diseases. 2001 Mar;32(6):897–928.

5. Dantas-Torres F, Chomel BB, Otranto D. Ticks and tick-borne diseases: A One Health perspective. Trends in Parasitology. 2012 Oct;28(10):437–446.

6. Schwartz AM, Hinckley AF, Mead PS, Hook SA, Kugeler KJ. Surveillance for Lyme Disease — United States, 2008–2015. MMWR Surveillance Summaries. 2017 Nov;66(22):1–12.

7. Olds CL, Mason KL, Scoles GA. *Rhipicephalus appendiculatus* ticks transmit Theileria parva from persistently infected cattle in the absence of detectable parasitemia: implications for East Coast fever epidemiology. Parasites & Vectors. 2018 Dec;11(1):126.

8. Vanwambeke SO, Šumilo D, Bormane A, Lambin EF, Randolph SE. Landscape Predictors of Tick-Borne Encephalitis in Latvia: Land Cover, Land Use, and Land Ownership. Vector-Borne and Zoonotic Diseases. 2010 Jun;10(5):497–506.

9. Dantas-Torres F. Climate change, biodiversity, ticks and tick-borne diseases: The butterfly effect. International Journal for Parasitology: Parasites and Wildlife. 2015 Dec;4(3):452–461.

10. Lambin EF, Tran A, Vanwambeke SO, Linard C, Soti V. Pathogenic landscapes: Interactions between land, people, disease vectors, and their animal hosts. International Journal of Health Geographics. 2010 Oct;9(1):54.

11. Kimanzi JK, Wishitemi BEL. Effects of land use changes on herbivores of Masai Mara ecosystem. International Journal of Environmental Studies. 2001 Oct;58(6):727–740.

12. Serneels S, Said MY, Lambin EF. Land cover changes around a major east African wildlife reserve: The Mara Ecosystem (Kenya). International Journal of Remote Sensing. 2001 Jan;22(17):3397–420.

13. Mundia CN, Murayama Y. Analysis of Land use/cover changes and animal population dynamics in a wildlife sanctuary in East Africa. Remote Sensing. 2009 Nov;1(4):952–970.

14. The Maasai Mara Science and Development Initiative. Maasai Mara: The challenges of a world unique ecosystem. 2015. http://maasaimarascience.org/fileadmin/projects/masaimara/MMSDI_Policy_Paper_Final.pdf. Accessed 20 July 2019.

15. Lamprey RH, Reid RS. Expansion of human settlement in Kenya’s Maasai Mara: what future for pastoralism and wildlife? Journal of Biogeography. 2004 May;31(6):997–1032.

16. Ogutu JO, Owen-Smith N, Piepho H-P, Said MY. Continuing wildlife population declines and range contraction in the Mara region of Kenya during 1977-2009. Journal of Zoology. 2011 Oct;285(2):99–109.

17. Gakuya F, Ombui J, Heukelbach J, Maingi N, Muchemi G, Ogara W, et al. Knowledge of Mange among Masai Pastoralists in Kenya. Nishiura H, editor. PLoS ONE. 2012 Aug;7(8):e43342.

18. Ginsberg HS, Ewing CP. Comparison of flagging, walking, trapping, and collecting from hosts as sampling methods for northern deer ticks, *Ixodes dammini*, and lone-star ticks, *Amblyomma americanum* (Acari:Ixodidae). Experimental & applied acarology. 1989 Sep;7(4):313–22.

19. Walker A, Bouattour A, Camicas J, Estrada-Pena A, Horak I, Latif AA, et al. Ticks of domestic animals in Africa: a guide to identification of species. Edinburgh: Bioscience Reports; 2003.

20. Crowder CD, Rounds MA, Phillipson CA, Picuri JM, Matthews HE, Halverson J, et al. Extraction of total nucleic acids from ticks for the detection of bacterial and viral pathogens. Journal of medical entomology. 2010 Jan;47(1):89–94.

21. Mwamuye MM, Kariuki E, Omondi D, Kabii J, Odongo D, Masiga D, et al. Novel *Rickettsia* and emergent tick-borne pathogens: A molecular survey of ticks and tick-borne pathogens in Shimba Hills National Reserve, Kenya. Ticks and Tick-borne Diseases. 2017 Feb;8(2):208–218.

22. Fard SN, Khalili M. PCR-Detection of *Coxiella burnetii* in ticks collected from sheep and goats in Southeast Iran. Iranian journal of arthropod-borne diseases. 2011;5(1):1–6.

23. Omondi D, Masiga DK, Ajamma YU, Fielding BC, Njoroge L, Villinger J. Unraveling Host-Vector-Arbovirus Interactions by Two-Gene High Resolution Melting Mosquito Bloodmeal Analysis in a Kenyan Wildlife-Livestock Interface. PLoS One. 2015;10(7):e0134375.

24. Ogola E, Villinger J, Mabuka D, Omondi D, Orindi B, Mutunga J, et al. Composition of *Anopheles* mosquitoes, their blood-meal hosts, and *Plasmodium falciparum* infection rates in three islands with disparate bed net coverage in Lake Victoria, Kenya. Malaria journal. 2017;16(1):360.

25. Kearse M, Moir R, Wilson A, Stones-Havas S, Cheung M, Sturrock S, et al. Geneious Basic: An integrated and extendable desktop software platform for the organization and analysis of sequence data. Bioinformatics. 2012 Jun;28(12):1647–1649.

26. Guindon S, Dufayard J-F, Lefort V, Anisimova M, Hordijk W, Gascuel O. New Algorithms and Methods to Estimate Maximum-Likelihood Phylogenies: Assessing the Performance of PhyML 3.0. Systematic Biology. 2010 Mar;59(3):307–321.

27. Lefort V, Longueville J-E, Gascuel O. SMS: Smart Model Selection in PhyML. Molecular Biology and Evolution. 2017 Sep;34(9):2422–2424.

28. Lopez-Velez R, Palomar AM, Oteo JA, Norman FF, Pérez-Molina JA, Portillo A. Novel *Candidatus* rickettsia species detected in nostril tick from human, Gabon, 2014. Emerging infectious diseases. 2015 Feb;21(2):325–7.

29. Macaluso KR, Davis J, Alam U, Korman A, Rutherford JS, Rosenberg R, et al. Spotted fever group rickettsiae in ticks from the Masai Mara region of Kenya. The American journal of tropical medicine and hygiene. 2003 May;68(5):551–3.

30. Ndeereh D, Muchemi G, Thaiyah A, Otiende M, Angelone-Alasaad S, Jowers MJ. Molecular survey of *Coxiella burnetii* in wildlife and ticks at wildlife–livestock interfaces in Kenya. Experimental and Applied Acarology. 2017 Jul;72(3):277–289.

31. Lopes MG, Junior JM, Foster RJ, Harmsen BJ, Sanchez E, Martins TF, et al. Ticks and rickettsiae from wildlife in Belize, Central America. Parasites Vectors. 2016 Dec;9(1):62.

32. Lado P, Nava S, Mendoza-Uribe L, Caceres AG, Delgado-de la Mora J, Licona-Enriquez JD, Delgado-de la Mora D, Labruna MB, Durden LA, Allerdice ME, Paddock CD. The *Amblyomma maculatum* Koch, 1844 (Acari: Ixodidae) group of ticks: phenotypic plasticity or incipient speciation? Parasites & vectors. 2018 Dec;11(1):610.

33. Kimita G, Mutai B, Nyanjom SG, Wamunyokoli F, Waitumbi J. Phylogenetic variants of *Rickettsia africae*, and incidental identification of “*Candidatus* Rickettsia Moyalensis” in Kenya. PLOS Neglected Tropical Diseases. 2016 Jul;10(7):e0004788.

34. Koka H, Sang R, Kutima HL, Musila L, Macaluso K. The detection of spotted fever group rickettsia DNA in tick samples from pastoral communities in Kenya. Journal of Medical Entomology. 2017 May;54(3):774–780.

35. Raoult D, Fournier PE, Fenollar F, Jensenius M, Prioe T, De Pina JJ, et al. *Rickettsia africae*, a tick-borne pathogen in travelers to sub-Saharan Africa. New England Journal of Medicine. 2001 May;344(20):1504–1510.

36. Rutherford JS, Macaluso KR, Smith N, Zaki SR, Paddock CD, Davis J, et al. Fatal Spotted Fever Rickettsiosis, Kenya. Emerging Infectious Diseases. 2004;10(5):910–913.

37. Yoshikawa H, Kimura M, Ogawa M, Rolain J-M, Raoult D. Laboratory-confirmed Mediterranean spotted fever in a Japanese traveler to Kenya. The American journal of tropical medicine and hygiene. 2005 Dec;73(6):1086–9.

38. Thiga JW, Mutai BK, Eyako WK, Ng’ang’a Z, Jiang J, Richards AL, et al. High Seroprevalence of Antibodies against Spotted Fever and Scrub Typhus Bacteria in Patients with Febrile Illness, Kenya. Emerging Infectious Diseases. 2015 Apr;21(4):688–691.

39. Maina AN, Farris CM, Odhiambo A, Jiang J, Laktabai J, Armstrong J, et al. Q fever, scrub typhus, and rickettsial diseases in children, Kenya, 2011–2012. Emerging Infectious Diseases. 2016 May;22(5):883–886.

40. Omballa VO, Musyoka RN, Vittor AY, Wamburu KB, Wachira CM, Waiboci LW, et al. Serologic evidence of the geographic distribution of bacterial zoonotic agents in Kenya, 2007. American Journal of Tropical Medicine and Hygiene. 2016 Jan;94(1):43–51.

41. Parola P, Paddock CD, Socolovschi C, Labruna MB, Mediannikov O, Kernif T, et al. Update on Tick-Borne Rickettsioses around the World: A Geographic Approach. Clinical Microbiology Reviews. 2013 Oct 1;26(4):657–702.

42. Uchiyama T, Kawano H, Kusuhara Y. The major outer membrane protein rOmpB of spotted fever group rickettsiae functions in the rickettsial adherence to and invasion of Vero cells. Microbes and Infection. 2006 Mar;8(3):801–809.

43. Maina AN, Jiang J, Omulo SA, Cutler SJ, Ade F, Ogola E, et al. High prevalence of Rickettsia africae variants in *Amblyomma variegatum* ticks from domestic mammals in rural western Kenya: implications for human health. Vector borne and zoonotic diseases. 2014 Oct;14(10):693–702.

44. Norval RAI, Perry BD, Young AS. The epidemiology of theileriosis in Africa. London: Academic Press; 1992. 481 p. Available from: https://cgspace.cgiar.org/handle/10568/91064

45. Perry BD. The control of East Coast fever of cattle by live parasite vaccination: A science-to-impact narrative. One Health. 2016 Dec;2:103–114.

46. Magulu E, Kindoro F, Mwega E, Kimera S, Shirima G, Gwakisa P. Detection of carrier state and genetic diversity of *Theileria parva* in ECF-vaccinated and naturally exposed cattle in Tanzania. Veterinary Parasitology: Regional Studies and Reports. 2019 Aug;17.

47. Nijhof AM, Bodaan C, Postigo M, Nieuwenhuijs H, Opsteegh M, Franssen L, et al. Ticks and associated pathogens collected from domestic animals in the Netherlands. Vector-Borne and Zoonotic Diseases. 2007;7(4):585–596.

48. Zhong J. Coxiella-like endosymbionts. Advances in Experimental Medicine and Biology. 2012;984:365–379.

49. Omondi D, Masiga DK, Fielding BC, Kariuki E, Ajamma YU, Mwamuye MM, et al. Molecular detection of tick-borne pathogen diversities in ticks from livestock and reptiles along the shores and adjacent islands of Lake Victoria and Lake Baringo, Kenya. Frontiers in Veterinary Science. 2017 Jun;4:73.

50. Khoo JJ, Lim FS, Chen F, Phoon WH, Khor CS, Pike BL, et al. *Coxiella* detection in ticks from wildlife and livestock in Malaysia. Vector-Borne and Zoonotic Diseases. 2016;16(12):744–751.

51. Smith TA, Driscoll T, Gillespie JJ, Raghavan R. A *Coxiella*-like endosymbiont is a potential vitamin source for the Lone Star tick. Genome Biology and Evolution. 2015 Mar;7(3):831–838.

52. Duron O. The IS1111 insertion sequence used for detection of *Coxiella burnetii* is widespread in *Coxiella*-like endosymbionts of ticks. FEMS Microbiology Letters. 2015 Aug;362(17).

53. Jourdain E, Duron O, Barry S, González-Acuña D, Sidi-Boumedine K. Molecular methods routinely used to detect *Coxiella burnetii* in ticks cross-react with Coxiella-like bacteria. Infection Ecology & Epidemiology. 2015 Jan;5(1):29230.

54. Machado-Ferreira E, Vizzoni VF, Balsemão-Pires E, Moerbeck L, Gazeta GS, Piesman J, et al. *Coxiella* symbionts are widespread into hard ticks. Parasitology Research. 2016;115(12):4691–4699.

55. Randolph SE, Green RM, Hoodless AN, Peacey MF. An empirical quantitative framework for the seasonal population dynamics of the tick *Ixodes ricinus*. International journal for parasitology. 2002 Jul;32(8):979–89.

56. Sojka D, Franta Z, Horn M, Caffrey CR, Mareš M, Kopáček P. New insights into the machinery of blood digestion by ticks. Trends in Parasitology. 2013;29(6):276–285.

57. Brahma RK, Dixit V, Sangwan AK, Doley R. Identification and characterization of *Rhipicephalus (Boophilus) microplus* and *Haemaphysalis bispinosa* ticks (Acari: Ixodidae) of North East India by ITS2 and 16S rDNA sequences and morphological analysis. Exp Appl Acarol. 2014;62(2):253–65.

58. Hebert PDN, Penton EH, Burns JM, Janzen DH, Hallwachs W. Ten species in one: DNA barcoding reveals cryptic species in the neotropical skipper butterfly *Astraptes fulgerator*. Proc Natl Acad Sci U S A. 2004;101(41):14812–7.

59. Chitimia L, Lin RQ, Cosoroaba I, Braila P, Song HQ, Zhu XQ. Molecular characterization of hard and soft ticks from Romania by sequences of the internal transcribed spacers of ribosomal DNA. Parasitol Res. 2009;105(4):907–11.

60. Roux V, Raoult D. Phylogenetic analysis of members of the genus Rickettsia using the gene encoding the outer-membrane protein rOmpB (ompB). Int J Syst Evol Microbiol. 2000;50(4):1449–55.

61. Tokarz R, Kapoor V, Samuel JE, Bouyer DH, Briese T, Lipkin WI. Detection of tick-borne pathogens by masstag polymerase chain reaction. Vector-Borne Zoonotic Dis. 2009;9(2):147–51.

62. Georges K, Loria GR, Riili S, Greco A, Caracappa S, Jongejan F, et al. Detection of haemoparasites in cattle by reverse line blot hybridisation with a note on the distribution of ticks in Sicily. Vet Parasitol. 2001;99(4):273–86.

63. Hoover TA, Vodkin MH, Williams JC. A *Coxiella burnetii* repeated DNA element resembling a bacterial insertion sequence. J Bacteriol. 1992;174(17):5540–8.

64. Boakye DA, Tang J, Truc P, Merriweather A, Unnasch TR. Identification of bloodmeals in haematophagous Diptera by cytochrome B heteroduplex analysis. Med Vet Entomol. 1999;13(3):282–7.

